# Global genomic epidemiology of *bla*_GES-5_ carbapenemase-associated integrons

**DOI:** 10.1101/2024.02.09.579496

**Authors:** William Matlock, Liam P. Shaw, Nicole Stoesser

**Author notes:** **Correspondence:** William Matlock and Nicole Stoesser. Contributed equally as senior authors (Liam P. Shaw, Nicole Stoesser).

## Abstract

Antimicrobial resistance (AMR) gene cassettes comprise an AMR gene flanked by short recombination sites (*attI* × *attC* or *attC* × *attC*). Integrons are genetic elements able to capture, excise, and shuffle these cassettes, providing ‘adaptation on demand’, and can be found on both chromosomes and plasmids. Understanding the patterns of integron diversity may help to understand the epidemiology of AMR genes. As a case study, we examined the clinical resistance gene *bla*_GES-5_, an integron-associated class A carbapenemase first reported in Greece in 2004 and since observed worldwide, which to our knowledge has not been the subject of a previous global analysis. Using a dataset comprising all NCBI contigs containing *bla*_GES-5_ (*n* = 431), we developed a pangenome graph-based workflow to characterise and cluster the diversity of *bla*_GES-5_ -associated integrons. We demonstrate that *bla*_GES-5_-associated integrons on plasmids are different to those on chromosomes. Chromosomal integrons were almost all identified in *P. aeruginosa* ST235, with a consistent gene cassette content and order. We observed instances where insertion sequence IS*110* disrupted *attC* sites, which might immobilise the gene cassettes and explain the conserved integron structure despite the presence of *intI1* integrase promoters, which would typically facilitate capture or excision and rearrangement. The plasmid-associated integrons were more diverse in their gene cassette content and order, which could be an indication of greater integrase activity and ‘shuffling’ of integrons on plasmids.

## Introduction

The dissemination of antimicrobial resistance (AMR) genes combines multiple modes of transfer, operating within a hierarchy of genetic units: AMR genes, their immediate genetic contexts, mobile genetic elements (MGEs), the bacterial host, and microbial communities^1^, similar to the structure of a nested ’Russian-doll’^2^. The combined actions of these units mean the epidemiological picture of AMR gene dissemination can quickly become complex.

The immediate genetic context, or ‘flanking sequences’, of AMR genes can act as epidemiologically relevant markers. These sequences can contain a great deal of genetic diversity, encompassing an array of other genes, MGEs, and other structures, including integrons^3^. Structurally, class 1 integrons consist of an integrase (*intI1*) upstream of a series of genes. Each gene is flanked by recombination sites (*attI* × *attC* and *attC* × *attC* sites in the first and subsequent positions, respectively) and the series of genes form an array of gene ‘cassettes’ (Figure 1). The integron integrase belongs to the family of site-specific tyrosine recombinases, including phage integrases such as Frat1 and D29, and XerC and XerD in *E. coli*^4^. Whilst most genes in this family share the conserved residue RHRY, alongside other motifs (boxes I and II, patches I, II and III), integron integrases uniquely possess an additional ∼30 residues near the patch III motif ^5^, which is thought to aid the movement of gene cassettes^6^. This behaviour also enables the integron to vary the order of the gene cassettes, which in turn modulates their relative expression by proximity to the cassette promoter (P_c_; found within the integrase)^7^. This variable order permits ‘adaptation on demand’ in response to changing conditions, with the array of cassettes acting as a store of previously beneficial genes^8^. The expression of the integrase is controlled separately by its promoter (P_int_; within or downstream of the integrase). Class 1 integrons are found on both chromosomes and plasmids (known as ’mobile integrons’)^3^. On plasmids, the integron can be transferred between cells. Then, within the new cell, the gene cassettes can be captured by other integrons^9^. In this way, plasmids and integrons work in tandem to disseminate AMR genes.

**Figure 1.**
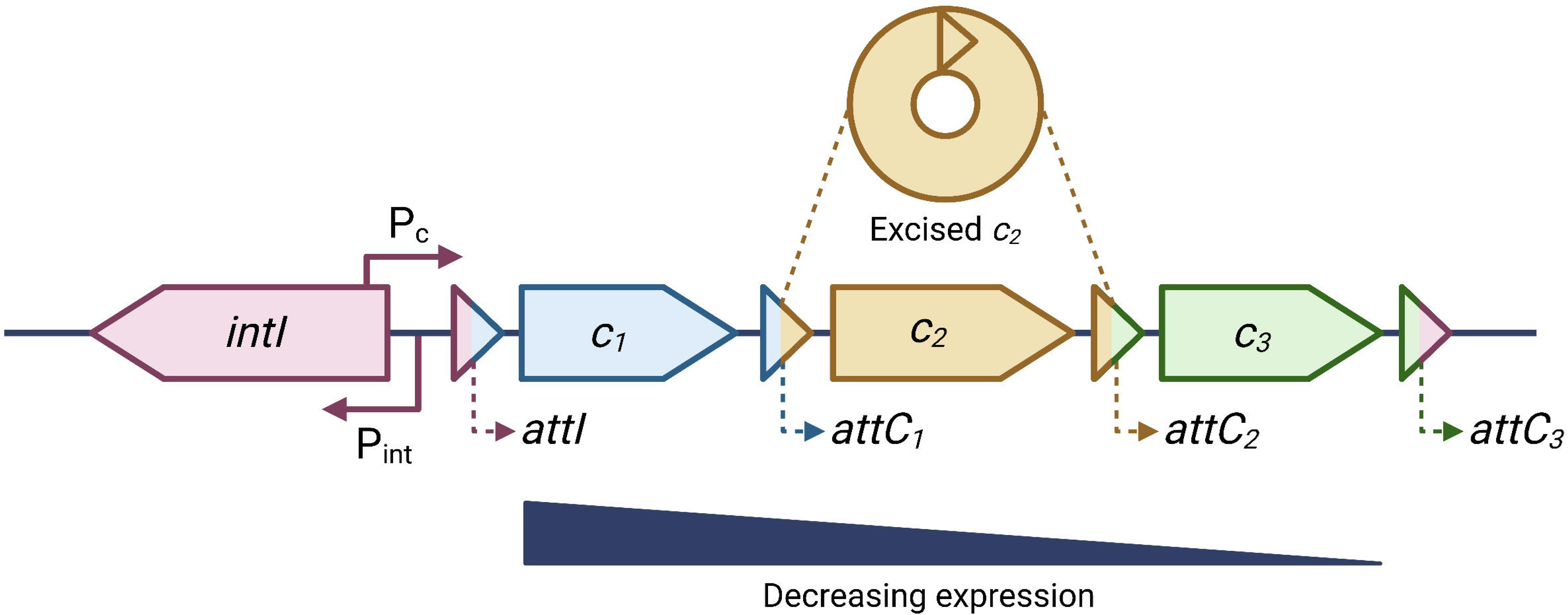
A schematic of an integron. Adapted from Gillings 2017^11^ and Ghaly 2017^11^. Shown is an integrase in the reverse orientation with a cassette promoter (P_c_) and integrase promoter (P_int_). This can lead to transcriptional interference^8^. Also shown are three gene cassettes (*c_1_*, *c_2_*, and *c_3_*), flanked by the attachment sites (*attI*, *attC_1_*, *attC_2_*, and *attC_3_*). Gene cassette c2 has been excised above the integron. The expression of the gene cassettes generally decreases with distance from the promoter P_c_.

Historically, class 1 integrons have been associated with AMR gene cassettes conferring resistance to several classes of antibiotics, including aminoglycosides (AAC/AAD-family resistant N-acetyltransferases), beta-lactams (GES/IMP/VIM-family beta-lactamases), trimethoprims (Dfr-family dihydrofolate reductases), and sulphonamides (Sul-family dihydropteroate synthases)^3^. Class 1 integrons are believed to have jumped from environmental to human species via a Tn*402* transposon^10,11^. Sulphonamides are the oldest widely used antibiotics (discovered in the mid-1930s), giving any clinical integrons with a *sul1* gene cassette a selective advantage. Alongside *sul1*, early clinical class 1 integrons also carried the *qacE* gene cassette, conferring resistance to biocides, and a mercury resistance *mer* Tn*501*-like transposon^11^. The structures of contemporary class 1 integrons are still associated with many of these structures downstream of the gene cassettes in a 3’ conserved sequence^12^, as well as the whole integron being flanked by the inverted repeats associated with Tn*402*^13^. However, class 1 integrons also carry a diversity of more recently evolved gene cassettes^14^. Currently, integron integrases are so prolific in human microbiota that they have been used as a proxy for anthropogenic pollution^15^. Class 2, 3, 4, and 5 integrons are also observed in both clinical and environmental niches, and associated with different integrases, niches, and gene cassette arrays^9,12,14^.

Currently, there are several software tools for the epidemiological study of AMR. These include those focussing on non-plasmid MGEs such as transposable elements^16,17^, as well as whole plasmids^18–20^ and bacterial strains that commonly carry particular AMR genes^21^. For integrons, epidemiology has often relied on sequence comparisons to reference databases, such as INTEGRALL^14^, so is restricted to known diversity. Moreover, due to the rearrangement of gene cassettes, integrons containing the same cassette array but in a different order can have poor pairwise alignment scores (for example, with MAFFT^22^), diminishing the utility of traditional alignment-based methods. Lastly, the repetitive structures of integrons make sequence assembly challenging, often generating short or ‘broken’ contigs^23–25^.

Alternatively, integron epidemiology can use phylogenetic-based approaches. Integron integrases are highly conserved genes^5,26^, with a strong phylogenetic signal for host genera and sampling environment^27–29^. This signal has similarly been demonstrated for *attC* site variants and is arguably connected: both the integrase and *attC* sites must cooperate well to successfully modulate the gene cassette array under selective pressure^30^. Previous studies have demonstrated the potential of tracking single nucleotide variants (SNVs) of the whole integron sequence across a single outbreak^31^, and synonymous single nucleotide variants (sSNVs) of individual gene cassettes across GenBank sequences^3^. Though synonymous mutations might leave the protein unchanged, they can still alter the fitness^32^, for instance, by increasing translational efficiency^33,34^. Within a bacterial population, tracking SNVs might indicate whether an integron has emerged locally or has been imported, and hence improve our understanding of how specific integrases and gene cassettes have moved historically. This is particularly important for mobile integrons. Overall, reference-free methods that are robust to both variable gene cassette synteny and ‘broken’ contigs would be a useful addition to studying integron and AMR gene epidemiology.

In this study, we describe a novel computational workflow for integron epidemiology. Capturing the diversity and synteny of gene cassettes in a sample of genomes has parallels to pangenome graph construction, where the aim is to summarise the sharing of genetic structures within whole genomes. Here, we leverage recent advances in pangenome graphs to cluster a large dataset of sequences containing a specific integron-associated AMR gene, *bla*_GES-5_. We show that this approach is concordant with genetic structure as well as bacterial niche and that it is robust to both variable gene cassette synteny and the inclusion of short or incomplete contigs. In addition, we examine the potential of SNV profiling for global integron epidemiology.

## Materials and methods

The GitHub repository https://github.com/wtmatlock/ges contains a tutorial for the scripts and commands used in this analysis.

### Distribution of NCBI GES variant annotations

All GES genes (*n*=57) were retrieved from the NCBI Reference Gene Catalog (as of 17/02/23). All alleles were 864bp except GES-42 (NG_065870.1:1-873) with a 9bp insertion. Protein sequences were aligned using clustalw^35^ (v 2.1; ‘slow’ mode), which was also used to produce a neighbour-joining tree, correcting for multiple substitutions. The hydrolytic profile was determined from experimental studies, collated by BLDB^36^ (as of 17/02/23). NCBI GES variant genera distribution was determined from querying NCBI’s ‘Pathogen detection <u>micro</u>bial <u>b</u>rowser for identification of <u>g</u>enetic and <u>g</u>enomic <u>e</u>lements’ (MicroBIGG-E) service^37^ using the query ‘element_symbol:blaGES*’ (as of 27/02/2023).

### Dataset curation

Using the same query as above, *n*=1,375 contigs were retrieved from NCBI’s (MicroBIGG-E service (as of 27/02/2023). These contigs were originally annotated using AMRFinderPlus^38^, a BLAST/HMMER based program reliant on the NCBI Pathogen detection reference gene catalogue. Contigs were then annotated for *bla*_GES-5_ using the NCBI GES-5 protein reference sequence WP_012658785.1 and BLASTX^39^. Filtering by 100% amino acid identity/coverage and a single hit yielded *n*=431 contigs.

NCBI BioProjects can contain multiple isolates from clonal strains and outbreaks, as well as from more general repeat sequencing. This meant there were potential biases in the *n*=431 *bla*_GES-5_-positive contigs, which could lead to the artificial inflation of some flanking structures and/or isolate metadata in the downstream analysis. To help correct this, the contigs were deduplicated as follows (see deduplication.R): first, the longest contig from each BioSample was kept and the rest discarded. If there were ties for the longest contig, a random representative was chosen, yet this was never the case. Then, within each BioProject, contigs that were perfectly contained in another were discarded, to diminish the bias of flanking sequence duplicates with the same metadata. Pairwise contig containment was calculated using Mash screen^40^ with code given in the repository. If this resulted in all contigs from a BioProject being discarded, we chose a random, longest representative. This left *n*=107 contigs. Finally, *n*=3 contigs were removed from the dataset because they contained repeated *bla*_GES-5_ PanGraph blocks in the 10kbp flanking sequences: adjacent blocks in NZ_CP073313.1 and NZ_VAAM01000031.1, and ∼9kbp apart in NZ_UARQ01000013.1 with several other PanGraph blocks repeated. Whilst these cases were potentially assembly errors, they also made pangenome graph construction challenging. This left a total of *n*=104 contigs for analysis.

Contig lengths were calculated with fastaLengths.py in the repository, which depends on Biopython^41^. BioProject accessions were retrieved using NCBI’s E-utilities^42^ via contig BioSample accessions. The code used is given in the repository README.md tutorial. Pairwise sequence containment used Mash screen^40^ (v. 2.2; default parameters except sketch size -s 100000).

### Exploring *bla*_GES-5_ flanking sequences with Flanker

Flanking sequences were extracted and clustered using the Python tool Flanker with a custom Abricate (v. 1.0.1; Seeman 2020) database containing only the NCBI *bla*_GES-5_ nucleotide reference sequence NG_049137.1. The Flanker command used was flanker --gene GES-5 -- database GES-5 --fasta_file ./“$f”.fasta --flank both --window 10000 --wstop 10000 --wstep 100 --include_gene -cl. Flanker was run by the Slurm scheduler script runFlanker.sh, but could be adapted to any command line.

The Flanker output was then processed (see plotFlanker.sh). First, matrices were generated for both the upstream and downstream flanks, where each row represented a contig, and each column the flanking sequence length (100bp, 200bp, …, 10,000bp). Each entry was either (i) a unique Flanker cluster label or (ii) 9999 if the flanking sequence had no cluster label because it was too short. Then, the row-wise Jaccard index was calculated to generate a distance matrix, which was hierarchically clustered. This generated the ‘meta-clusters’ (a clustering of the Flanker cluster profiles). *n*=20/104 and 7/104 sequences were removed upstream and downstream, respectively, because they had insufficient flanking sequence to be clustered.

### Sequence annotations

Contig annotations were pre-generated by NCBI’s Prokaryotic Genome Annotation Pipeline^43^, and retrieved via contig accessions. The code used to download the GFF files is given in the repository README.md tutorial. Additional annotations of integron integrases, *attC*/*attI* sites, and integron promoters P_c_/P_int_ were generated using IntegronFinder^44^ (v. 2.0.2; default parameters except --local-max --promoter-attI –lin; see runIntegronFinder.sh). Integron finder annotates integrases using hidden Markov models protein profiles, and *attC* sites using covariance models to predict secondary structure. For *attI* sites and promoters, reference motifs are searched for in query sequences. Integrase type (*intI1* or *intI3*) was confirmed using BLASTX^39^. Lastly, transposase additions were generated with Abricate^45^ (v. 1.0.1) with the ISfinder database^46^ (as of 28/02/2023) using default parameters except --db isfinder (see runISfinder.sh).

### Clustering *bla*_GES-5_ flanking sequences with PanGraph

The 10kbp *bla*_GES-5_ flanking sequences (extracted with Flanker as described above) were passed into PanGraph^47^ (v. 0.6.3) with pangraph build, then pangraph export --edge- minimum-length. BLASTN^48^ (v. 2.5.0) determined which of the PanGraph blocks contained NG_049137 (see runPanGraph.sh). An additional post-processing script converted the PanGraph GFA output to a visualization of coloured linear blocks (see pangraphGFA.py).

### Determining contig origin

Contigs were classified as chromosomal if they could be typed with mlst^49^ (v. 2.23.0) using default parameters (see runMlst.sh). Contigs were classified as plasmid if they could be typed with either Abricate^45^ (v. 1.0.1) and the PlasmidFinder database^18^ (as of 28/02/2023) using default parameters except --db plasmidfinder (see runPlasmidFinder.sh), or with MOB-Typer^50^ (v. 3.1.4) using default parameters (see runMobTyper.sh). Contigs were left unclassified if none of these methods gave a typing.

### SNV profiling

First, the integrase annotation positions were extracted from the IntegronFinder output. Then, the integrase sequences were retrieved from the full contigs using extractIntegrase.py in the repository. Integrase sequences were aligned using ClustalW^35^ (v. 2.1) on slow/accurate mode. The SNP distance matrix was generated using snp-dists^51^ (v. 0.8.2) with default parameters, and SNP constant sites were found using SNP-sites^52^ (v. 2.5.1) with -C and otherwise default parameters. The SNP clustering/heatmap was generated using the script plotHeatmap.R. The method was identical for *bla*_GES-5_, except the sequences were instead extracted using runFlanker.sh but altering the window to 0bp with --window 0.

### Data visualisation

Figures 2, S1, and S3-5 were generated using ggplot2^53^ in R. Figure 1 was drawn using BioRender.com. Figure S2 was produced using Bandage^54^. Figure 3 was plotted using plotFlanker.R. For plotting Figure 4, as well as statistics for contigs/annotations/PanGraph, we used plotFlanks.R.

**Figure 2.**
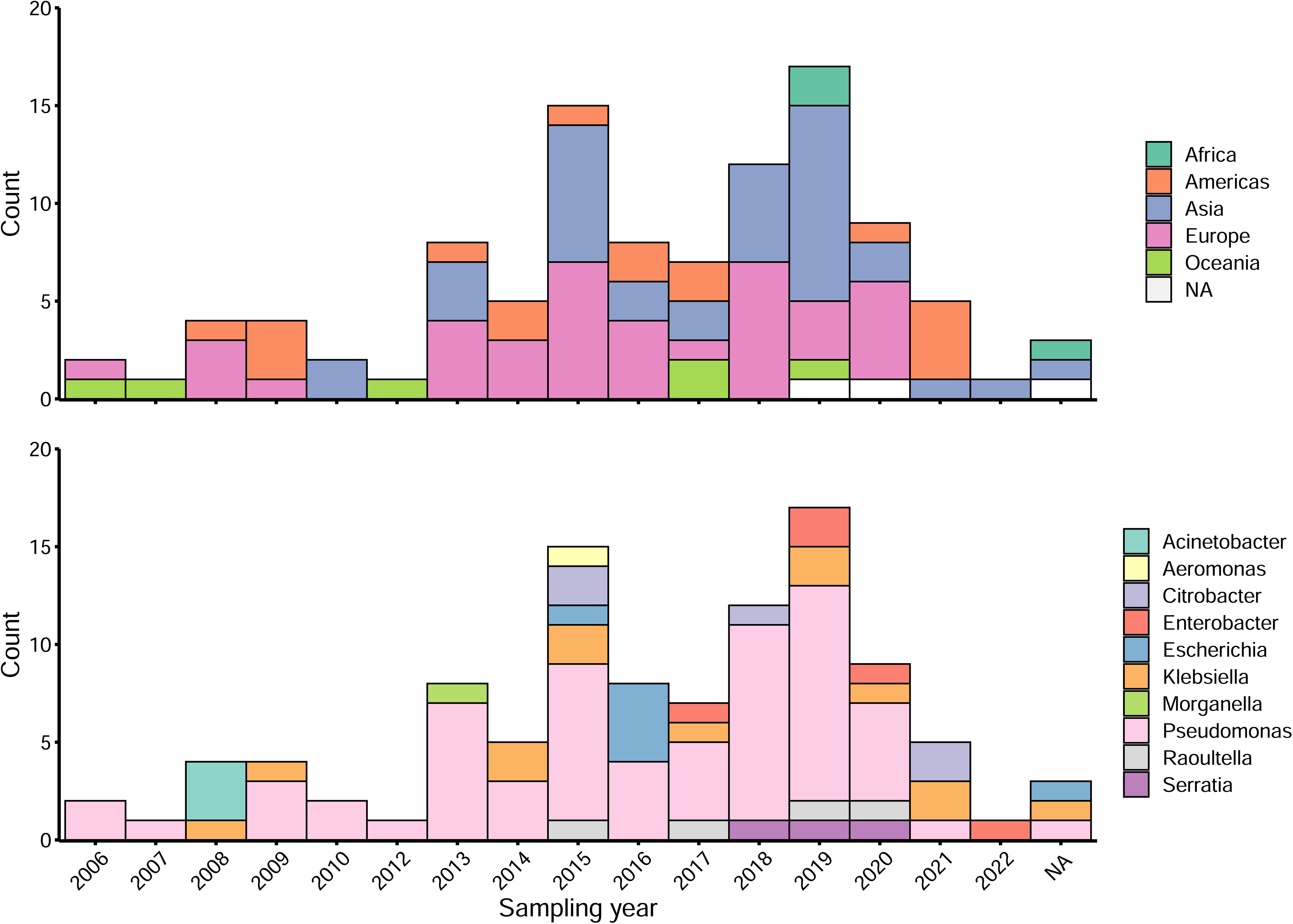
NCBI metadata distributions for the *n*=104 *bla*_GES-5_ positive contigs. **(a)** Shows ISO region and **(b)** isolate genus by sampling year. Note, no contigs were sampled in 2011 so this year is omitted from the *x*-axis.

**Figure 3.**
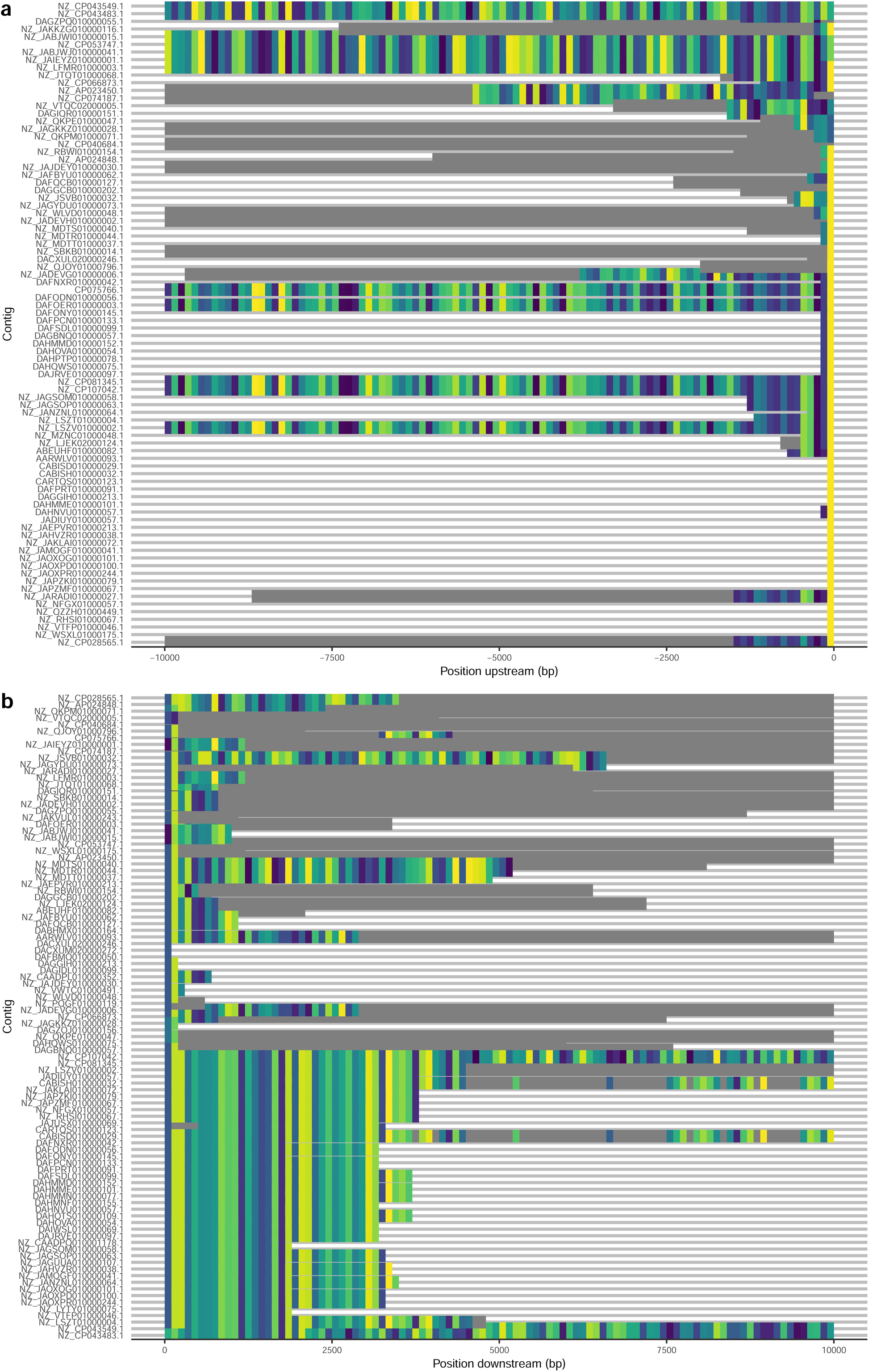
Exploration of flanking sequence diversity. **(a)** Upstream Flanker output, from 100bp flanking sequences to 10,000bp flanking sequences in 100bp windows. For each length, tile colours represent clusters found multiple times, and grey values represent unique clusters. Colours do not compare across lengths, only within lengths. **(b)** Downstream Flanker output, as described for the upstream flanking sequences.

**Figure 4.**
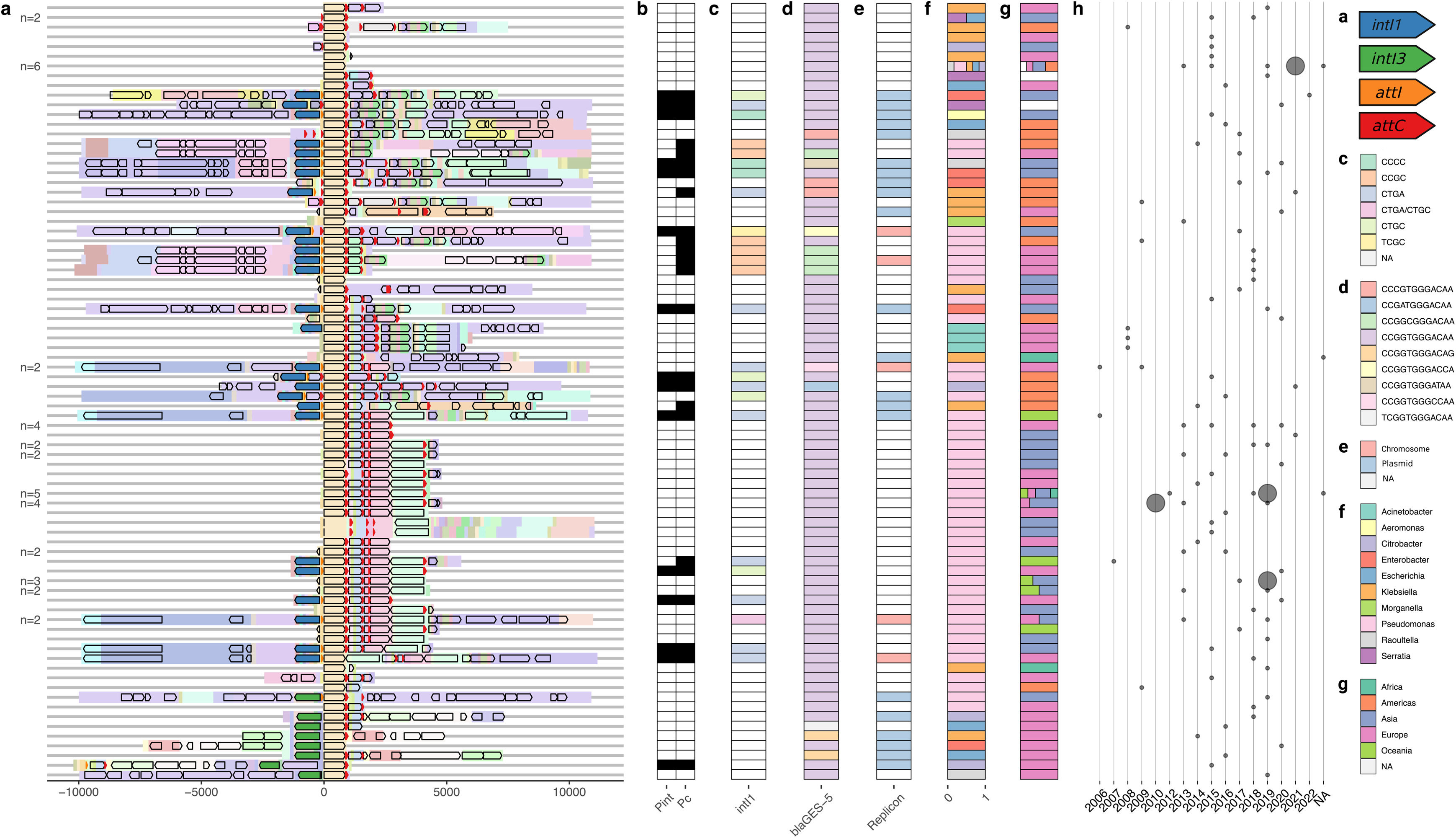
10,000bp flanking sequence analysis for bla_GES-5_. **(a)** Clustering-ordered PanGraph blocks (pastel colours) overlayed with CDS annotations (black outlined arrows), and integron-specific annotations: *intI1* in blue, *intI3* in green, *attI* in orange, and *attC* in red. Numbers to the left indicate duplicates. **(b)** P_int_ and P_c_ presence (black) or absence (white). **(c)** *intI1* SNV profiles, corresponding to the blue *intI1* annotations in panel (a). **(d)** *bla*_GES-5_ SNV profiles. **(e)** Contig origin (chromosomal, plasmid, or NA). **(f)** Contig genus distribution. **(g)** Contig ISO region distribution. **(h)** Contig sampling year distribution. Smaller circles indicate 1 isolate, and larger circles indicate 2 isolates.

## Results

### A historical and global dataset of *bla*_GES-5_ contigs

Public sequence databases such as NCBI present a valuable resource for the study of AMR gene epidemiology, despite potential sampling biases. The historic and continued deposition of sequence data means exploratory studies can represent a global epidemiological picture. For example, a recent study leveraged NCBI to analyse over 7,000 *bla*_NDM_-positive contigs, demonstrating the joint role of Tn*125*, Tn*3000*, and IS*26* in the gene’s global spread^55^.

The GES family of class A beta-lactamases contains *n*=57 protein variants as of July 2023^36^, the first of which, GES-1, was reported in 1999 on a 140kb plasmid in a *Klebsiella pneumoniae* ^56^. GES variants are usually found as class 1/3 integron gene cassettes^57^ and many demonstrate carbapenemase activity^36^. This analysis focuses on *bla*_GES-5_ since it is both a carbapenemase and the most abundant GES enzyme annotated in NCBI sequence data (Figure S1; see Materials and Methods). To date, *bla*_GES-5_ has been linked to several clinical outbreaks in Japan^58^, Korea^59^, the UK^60^, and the Czech Republic^61^. Moreover, *bla*_GES-5_ has been identified globally across a range of species, both on the chromosome (most notably *Pseudomonas aeruginosa* ST235^62^) and plasmids (for example, the Q-type plasmid pHPRU111 found across several *Enterobacterales spp*.^60^). It is also possible the abundance of GES-5 and other GES variants is underestimated due to a lack of targeted assays^63,64^.

We first downloaded all *n*=1,375 contigs annotated by NCBI with a GES variant (see Figure S1). We then annotated these for *bla*_GES-5_, giving *n*=431 contigs. Finally, we then deduplicated to *n*=107 to control for outbreaks/isolate repeat sequencing (see Materials and Methods). We also removed *n*=3 contigs which we found complicated the subsequent pangenome graph construction (see Materials and Methods), leaving a total of *n*=104 contigs (see Table S1). Our deduplicated dataset spanned 2006-2022 (*n*=3 missing data), represented all ISO (International Organisation for Standardization) regions (*n*=3 Africa, *n*=17 Americas, *n*=36 Asia, *n*=39 Europe, *n*=6 Oceania, *n*=3 missing data; Figure 2a), and *n*=10 bacterial genera, of which n=63/104 (61%) sequences were from *Pseudomonas* and *n*=41/104 (47%) from other Gram-negative bacilli (*Acinetobacter* [*n*=3], *Aeromonas* [*n*=1], *Citrobacter* [*n*=5], *Enterobacter* [*n*=5], *Escherichia* [*n*=6], *Klebsiella* [*n*=13], *Morganella* [*n*=1], *Raoultella* [*n*=4], *Serratia* [*n*=3]; Figure 2b). The overrepresentation of *Pseudomonas* spp. is consistent with previously reported *bla*_GES-5_ distribution^62^. Assembled contigs containing *bla*_GES-5_ varied dramatically in length (median=4,900bp, range=960-7,086,823bp), reflecting the fact that assembly of integron regions from short reads can lead to short contigs.

### Exploring *bla*_GES-5_ flanking sequence diversity with Flanker

We first used Flanker^65^ for an exploratory coarse-grained clustering of the 10kbp flanking sequences both upstream and downstream of *bla*_GES-5_ (see Materials and Methods). Briefly, Flanker annotated the *n*=104 sequences for *bla*_GES-5_, then extracted flanking sequences in increasing 100bp steps in both the upstream and downstream directions, up to 10,000bp. Next, for each set of flanking sequences of a given length and direction, Flanker clustered them using Mash distances and single-linkage clustering. Figure 3 visualises these clusters for each contig. In addition to Flanker, we performed a secondary clustering of the cluster labels themselves (‘meta-clusters’; see Materials and Methods). In short, for both the upstream and downstream directions, we generated a matrix of contig cluster labels, calculated the Jaccard Index (*JI*) of these labels (the number of shared cluster labels divided by the total number of cluster labels; see Materials and Methods), and performed a complete hierarchal clustering. This grouped contigs with similar Flanker cluster profiles, revealing distinct meta-clusters both upstream and downstream of *bla*_GES-5_, consistent with shared genetic structures in the flanking sequences. The ordering of the hierarchal clustering dendrograms provided the order of the contigs in Figure 3.

### The genetic landscape of *bla*_GES-5_ integrons

Following the exploratory analysis with Flanker, we wanted to document the genetic structure of *bla*_GES-5_-associated integrons. To begin, we extracted flanking sequences 10kbp upstream and downstream of the gene (using Flanker, see Materials and Methods). For all the sequences, we retrieved annotation data from NCBI’s Prokaryotic Genome Annotation Pipeline (*n*=2/104 contigs had no annotations; see Materials and Methods). We also generated integron-specific annotations using Integron Finder and transposase annotations using the ISfinder database (see Materials and Methods). Using IntegronFinder, we annotated for (i) integrases, (ii) *attI* and *attC* sites, (iii) integrase promoters (P_int_), and (iv) cassette promoters (P_c_). In total, *n*=37 flanking sequences were annotated for an integrase, and of these, 73% (27/37) had an *attI* site and at least one *attC* site, indicating a complete integron. Notably, 22% (8/37) of integrase-positive sequences had at least one *attC* site but lacked an *attI* site, potentially consistent with the difficulty of annotating *attI* outside of class 1 integrons^44^. To predict putatively functional integrases, we included annotations for the promoters P_int_ and P_c_. Of the *n*=37 integrases annotated, 41% (15/37) had both P_int_ and P_c_, 24% (9/37) had just P_c_, and the remaining 35% (13/37) had neither, consistent with a spectrum of putative activity.

Using annotations found between either *attI* × *attC* or an *attC* × *attC* sites, we recovered *n*=39 putative unique gene cassettes and *n*=195 total cassettes across the flanking sequences (Table 1). For describing integrons in this study, gene cassettes are recorded between vertical bars ‘|’, with commas separating their genes e.g. |*a*|, |*b*, *c*| represents an integron array of two gene cassettes, one containing gene *a*, and another containing genes *b* and *c*. Gene cassettes were median=1 gene long (range=1-4 genes). *bla*_GES-5_ was almost always found alone as a gene cassette (34/35), except once with an IS*Pa21* insertion sequence and *aac(6’)-Ib4* (1/35). Aminoglycoside resistance genes were found in 51% (99/195) of gene cassettes, of which most were a lone *aac(6’)-Ib4* gene (42% [42/99]). Also observed were *dfrA*-family trimethoprim resistance genes (6/195), OXA-family beta-lactamase genes (3/195) and *ere(A)* macrolide resistance genes (2/195). Equally as common as the *aac(6’)-Ib4* cassette was a DUF1010 gene cassette (42/195), encoding an uncharacterised protein.

**Table 1.**
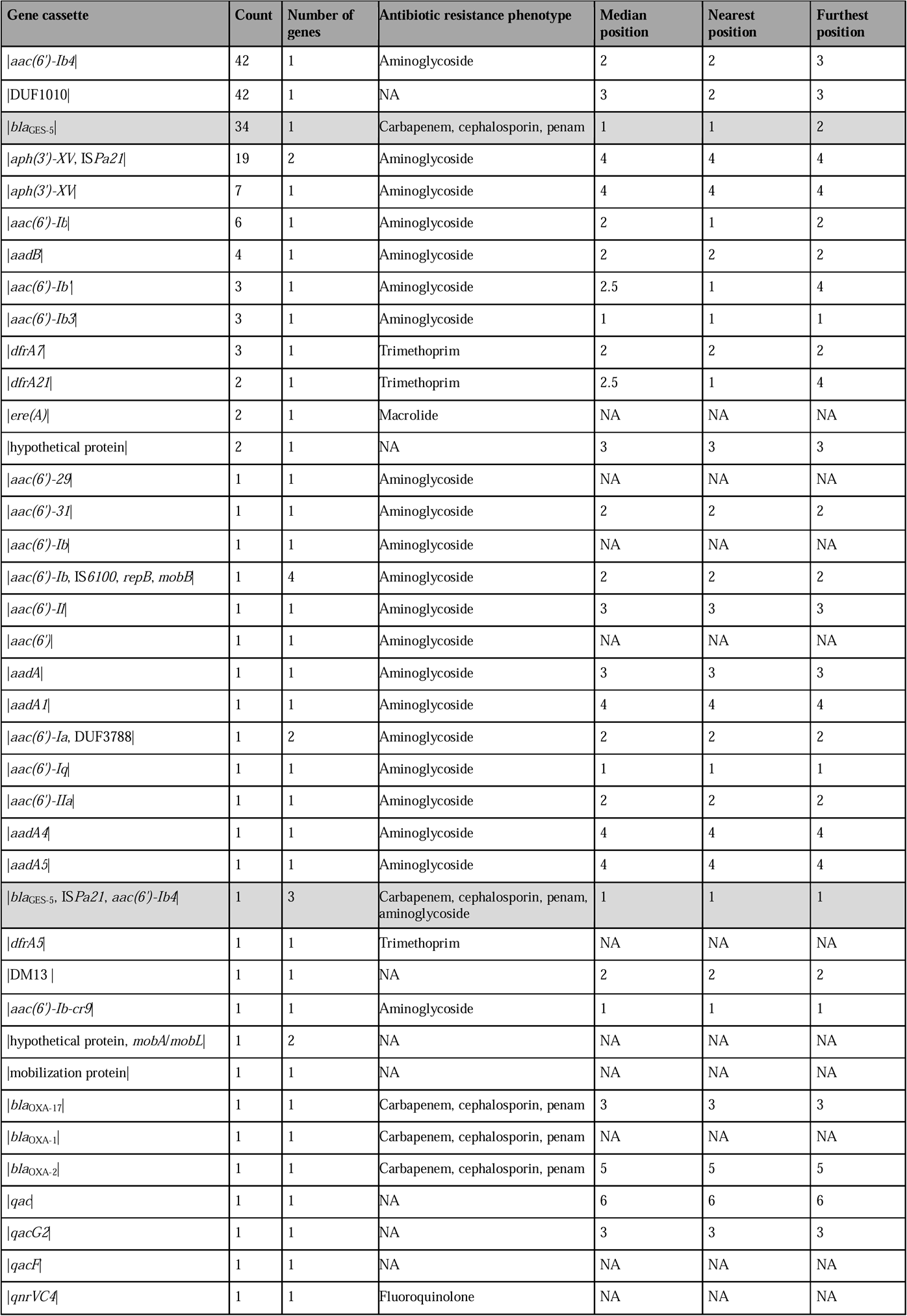
*Gene cassettes found on* bla*_GES-5_ integrons.* From left to right, the columns report the gene cassette, its count in the dataset, its length, its predicted antimicrobial resistance (AMR) phenotype, and its median, nearest, and furthest position from the integrase. Am., Ca., Ce, Fl., Ma., Pe., and Tr., denote predicted aminoglycoside, carbapenem, cephalosporin, fluoroquinolone, macrolide, penam, and trimethoprim resistance, respectively. When a gene cassette position could not be ascertained (for example, because it was found without an integrase upstream), NA is recorded. ‘hyp. prot.’ and ‘mob. prot.’ indicate hypothetical protein and mobility protein, respectively. Rows coloured grey indicate those containing *bla*_GES-5_.

In terms of insertion sequences, IS*Pa21* (of the IS*110* family, IS*1111* group) was observed *n*=18 and *n*=1 times in an |*aph(3’)-XV*, IS*Pa21*| and |*bla*_GES-5_, IS*Pa21*, *aac(6’)-Ib4*| gene cassette, respectively. The IS*1111* group is well-documented as targeting *attC* sites in class 1 integrons^66,67^. There was also *n*=1 instance of IS*1600* (of the IS*21* family) in an |*aac(6’)-Ib*, IS*6100*, *repB*, *mobB*| gene cassette. However, the terminal attachment site overlapped with a putative mobility protein, so was potentially a false positive.

We also examined the position of the gene cassettes downstream of the integrase (Table 1). To do this, we took the *n*=30/104 contigs with an integrase and/or *attI* site, and at least one gene cassette (as described above). Then, we recorded the position of the gene cassettes downstream of the integrase and *attI* site. The position of *n*=10/40 gene cassettes could not be determined as the contig(s) contained no integrase/*attI* site. Notably, *bla*_GES-5_ gene cassettes were almost always in the first position (93% [26/28]; where possible to calculate), consistent with the highest expression^68,69^. Overall, this approach is likely to underestimate the true number of gene cassettes in the sample due to difficulty annotating *attI* and *attC* sites, as well as fragmented contigs.

### Clustering *bla*_GES-5_ flanking sequences with PanGraph

Whilst the Flanker analysis captured some genetic similarity between the flanking sequences, it is order dependent: it can only compare sequence windows found in the same position upstream or downstream. In contrast, pangenome graph-based methods summarise the sharing of genes or genetic structures within a collection of sequences, independent of order. For the *bla*_GES-5_ flanking sequences, we generated a pangenome graph using PanGraph^47^ (Figure S2; see Materials and Methods). PanGraph represents each flanking sequence as a path through a graph where each node is an annotation-free multiple-sequence alignment (or ‘block’), and each edge is an inferred homology breakpoint. Stretches of sequence are collapsed into blocks by evaluating the costs of both within-block diversity and splitting a block. From the *n*=104 10kbp *bla*_GES-5_ flanking sequences, we extracted *n*=164 unique homology blocks, with median length=622.5bp (range=99-9,247bp). The graph totalled 247,747bp in length, of which 62% (153,992/247,747bp) comprised unique blocks (seen only in a single flanking sequence), consistent with substantial genetic diversity in *bla*_GES-5_ flanking sequences. The *n*=104 flanking sequences were represented by *n*=80 unique paths in the pangenome graph (median=1 flank per unique path, range=1-6).

Next, we clustered the flanking sequences by their putative gene cassette contents. First, for the *n*=69/80 flanking sequences containing either an *attI* and at least one *attC* site, or at least two *attC* sites, we collated a list of PanGraph blocks found between the maximally upstream and downstream sites. Then, for all the flanking sequences, we generated a presence/absence matrix of these blocks. This aimed to capture the gene cassette contents as blocks, even when *attI*/*attC* sites were poorly annotated, as well intergenic sequence. Using the presence/absence matrix, we calculated the pairwise *JI* of blocks between flanking sequences. Importantly, *JI* is indifferent to the ordering of blocks, such as those corresponding to shuffled gene cassettes from identical integrons. The resulting distance matrix was used for complete-linkage hierarchical clustering to order the flanking sequences (Figure 4a; Figure S3 presents a zoomed-in view for only the putative integron sequences, including annotations for the gene cassettes). We chose complete-linkage hierarchal clustering as it is robust to cluster shape, which we *a priori* did not know, and is less sensitive to outliers (for example, contigs with unique blocks) than other clustering approaches.

### The lifestyles of *bla*_GES-5_ integrons

Integrons can be found on both chromosomes and plasmids, so they can have different ‘lifestyles’ of transmission: either vertically on the chromosome or horizontally on plasmids, potentially moving between species. The inactivation of chromosomal integron integrases can fix a resistance phenotype within a strain^70^. In contrast, for conjugative plasmids, their transfer into a new cell can trigger an SOS response, which has been shown to upregulate integrase activity, leading to gene cassette gain, loss, and shuffling^71^. The genomic context of an integron is therefore crucial for understanding its epidemiology and evolution. To explore the lifestyles of *bla*_GES-5_ integrons, we compared their genomic contexts and sampling geography and year to their genetic structure.

We first assigned the *bla*_GES-5_ contigs as chromosomal or plasmid if a sequence type (ST) or replicon type could be identified (Figure 4e; see Materials and Methods). In total, 7% (7/107) contigs were classified as chromosomal (all *P. aeruginosa; n*=5 ST235, *n*=1 ST316, *n*=1 ST654), 20% (21/104) plasmid (*n*=5 IncP6, *n*=5 IncQ2, *n*=3 IncQ1, *n*=1 IncL/M, *n*=7 from untypeable replicon clusters) and 73% (76/104) neither, reflecting the short length of most of the contigs. Of the plasmids, 71% (15/21) were predicted to be mobilisable (19% [4/21] non-mobilisable, 10% [2/21] conjugative). In total, *n*=6 contigs were assigned both a ST and contained putative plasmid replicons (all *P. aeruginosa*, *n*=5 ST235, *n*=1 ST316). For ST235, *n*=5/5 contained an IncP replicon, as well as ‘rep_cluster_10’ and ‘rep_cluster_80’. One ST235 contig also contained “rep_cluster_322’. This is consistent with the identification of previously described genomic islands in ST235^72^. In addition to classifying the contigs by putative origin, we also integrated the NCBI sampling metadata from earlier: isolate genus (Figure 4f), sample ISO region (Figure 4g), and sampling year (Figure 4h; see Materials and Methods).

The PanGraph-based clustering revealed a family of *n*=42 class 1 integrons, of which *n*=2/42 were from *P. aeruginosa* ST235, n=1/42 contained ‘rep_cluster_80’, and n=39/42 were from unclassified contigs. The integron demonstrated a well-conserved gene cassette array: whilst only *n*=2/42 carried |*bla*_GES-5_|*aac(6’)-Ib4*|DUF1010|*aph(3’)−XV* |IS*Pa21*| with complete *attC* and *attI* boundaries, a total of *n*=30/42 carried these genes in the same order. Moreover, *n*=6/42 terminated before *ISPa21* with an *attC* site, and *n*=3/42 without. Most commonly, *ISPa21* inserted within an |*aph(3’)−XV*| cassette to form an |*aph(3’)−XV* , IS*Pa21*| cassette (16/42), but also once between |*bla*_GES-5_| and |*aac(6’)-Ib4*| to form a |*bla*_GES-5_, IS*Pa21*, *aac(6’)- Ib4*| cassette (1/42). Of the annotated promoters, 50% (5/10) had both P_int_ and P_c_, 20% (2/10) had no P_int_ or P_c_, and 10% (1/10) had P_c_ but no P_int_.

In contrast to the ST235-associated integrons, the *n*=21 plasmid-associated integrons were more varied, representing both *intI1* (10/21) and *intI3* (5/21; 6/21 broke off before the integrase). Within plasmid families, integron contents varied dramatically. Taking the largest identified replicon family, *n*=5 IncP6 plasmids, and gene cassettes with complete *attI*/*attC* boundaries, *n*=1/5 contained |*bla*_GES-5_|, *n*=1/5 |*bla*_GES-5_|, |*aac(6’)-31*|, |*qacG2*|, |*aadA5*|, *n*=1/5 |*bla*_GES-5_|, |*aac(6’)-IIa*|, |*aadA*|, *n*=1/5 |*bla*_GES-5_|, |*aac(6’)-Ia*, DUF3788|, and *n*=1/5 |*bla*_GES-5_|, |*aac(6’)-Ib4*|, |*qacF*|. No plasmids of the same replicon type shared the same gene cassette array.

### Integron SNV profiles are unreliable epidemiological markers

SNV profiling of the integrase and gene cassettes might help track integrons. However, there are potential pitfalls. Integrases are known to commonly undergo indel events, which can inactivate the gene and fix the gene cassettes in place^29^. Moreover, integrons can undergo frequent changes in cassette array, for instance under intermittent antimicrobial exposure^69^, meaning it might not be possible to consistently track specific gene cassette-integrase pairings. Using the curated dataset of *bla*_GES-5_ integron sequences, we explored single nucleotide variant (SNV) profiling of both the integrase and *bla*_GES-5_ gene.

For the integrase genes, we produced multiple-sequence alignments of the *n*=37 integrase annotations found in the flanking sequences (see Materials and Methods). This revealed two distinct clusters of SNV profiles (Figure S4). Cluster 1 (C1) contained *n*=29 *intI1* integrases median=1 SNP apart (range=0-3 SNPs), and were all 1,014bp in length except two at 867bp and 924bp, consistent with minimal indel events. Cluster 2 (C2) contained *n*=8 *intI3* integrases median=1.5 SNPs apart (range=0-14 SNPs), and were more variable in length (*n*=1 816bp, *n*=1 897bp, *n*=1 990bp, *n*=2 1,017bp, *n*=3 1,041bp), consistent with a higher diversity of indels. The integrase gene (*intI1* or *intI3*) is shown by colour in Figure 4a (blue or green, respectively). Additionally, *attI* and *attC* sites are shown in orange and red, respectively, and for clarity, the presence/absence of promoters P_int_/P_c_ are indicated in Figure 4b (see Materials and Methods).

For C1, we removed the two shorter integrases, aligned the remainder, and determined the SNV sites (see Materials and Methods). This recovered *n*=5 SNV profiles of length=4 nucleotides (positions 87, 90, 95, and 115): *n*=12 CTGA, *n*=6 CCGC, *n*=5 CTGC, *n*=3 CCCC, *n*=1 TCGC. The comparison of C1 SNV profiles against flanking sequence structure is shown in Figure 4c. In one case, a duplicated flanking sequence represented two SNV profiles (CTGA/CTGC).

Integrase SNV profiles showed some concordance with the resistance profile and wider flanking structure (see Figure S5 for *intI1* SNV profiles against flank block contents). For example, *intI1* profile CCGC was always found downstream of a conserved structure, containing *merA* (mercuric reductase; *n*=5/6 sequences, *n*=1/6 times the sequence did not continue upstream of *intI1*), which were also the only instances of the *merA* operon in the dataset. Moreover, 100% (6/6) of profile CCGC had P_c_ but no P_int_. Downstream of profile CCGC, the overall gene cassette array was more varied: *n*=3/6 carried |*bla*_GES-5_|, |*aadB*|, *n*=2/6 carried |*bla*_GES-5_|, and *n*=1/6 |*bla*_GES-5_|, |*aadB*|, |*aac(6’)-Il*|, |*aadA4*|. Alternatively, *intI1* profile CCCC was found with a variety of flanking structures both upstream (*n*=2/3 included *repB* and the toxin-antitoxin genes *brnT*/*brnA*, *n*=1/3 included the restriction enzyme *eco57IR*) and downstream (gene cassette arrays: *n*=1/3 |*bla*_GES-5_|, |*aac(6’)-31*|, |*qacG2*|, |*aadA5*|, *n*=1/3 |*bla*_GES-5_|, *n*=1/3 |*bla*_GES-5_, *aac(6’)-IIa*, *aadA*|). However, 100% (3/3) of profile CCCC had both P_c_ and P_int_. No SNV profile was consistently found with the same upstream/downstream structure or gene cassette array. Moreover, the integrase SNV profile did not always predict promoter presence: for instance, profile CTGA had both P_c_ and P_int_ 58% (7/12) of the time, missing P_int_ 17% (2/12), and had neither 25% (3/12). P_int_ can fall downstream of the integrase^8^, potentially causing an inconsistency between the *intI1* SNV profile and promoter presence.

For *bla*_GES-5_, we similarly produced multiple-sequence alignments of the *n*=104 annotations (all were 864bp long and encoded identical proteins). This revealed *n*=9 synonymous SNV (sSNV) profiles of length=12 nucleotides (positions 15, 97, 126, 171, 177, 195, 292, 494, 504, 696, 759, and 837), of which one represented 85% (88/104) of sequences (*n*=88 CCGGTGGGACAA, *n*=4 CCGGCGGGACAA, *n*=4 CCCGTGGGACAA, *n*=2 CCGGTGGGACAG, *n*=2 CCGGTGGGCCA, *n*=1 CCGGTGGGACCA, *n*=1 CCGGTGGGATAA , *n*=1 TCGGTGGGACAA, *n*=1 CCGATGGGACAA). The comparison of *bla*_GES-5_ sSNV profiles against flanking sequence structure is shown in Figure 4d. The dominant sSNV profile CCGGTGGGACAA was found across flanking structures and integrase SNV profiles. For the minority sSNV profiles, it was challenging to evaluate their utility as epidemiological markers, however, none consistently represented the same flanking structures. Of possible interest was sSNV profile CCGGCGGGACAA, which was always associated with integrase SNV profile CCGC and *Pseudomonas* spp., with *n*=1/4 contigs assigned chromosomal and the remainder unclassifiable. Overall, across the *bla*_GES-5_- associated integrons, there are globally dominant *intI1* (CCGGTGGGACAA) *bla*_GES-5_ (CTGA) SNVs. This diminishes the epidemiological value of using these SNVs on a global scale. Moreover, if these common variants are present in local contexts, such as in hospital or community surveillance, it might also be limiting.

## Discussion

Class 1 integrons play a crucial role in global AMR spread, yet they can be challenging to analyse and track. Limitations in sequencing and assembly methods result in fragmented contigs, difficulties in annotating the attachment sites leave ambiguities in the context and structure of gene cassettes, and reliance on reference databases restricts epidemiological efforts to known diversity. In this study, we have explored a new approach to integron genomic epidemiology, exemplified by the integron-associated carbapenemase gene *bla*_GES-5_, originally identified in 2004 on a class 1 integron^73^. To our knowledge, this is the first review of *bla*_GES-5_ global genomic epidemiology using all public *bla*_GES-5_-containing contigs, and indeed of any GES variant.

We first curated a global dataset of 104 representative *bla*_GES-5_ contigs from NCBI, spanning more than 15 years. These contained 39 unique gene cassettes, arranged in 17 unique complete arrays (namely those which started with either an integrase and *attI* sites and contain at least one complete gene cassette, see Figure S3). Many more arrays were likely sampled, but we were limited by missing gene and *attI* and *attC* site annotations, as well as ‘broken’ contigs. At the time of writing, the INTEGRALL database held just 9 unique gene cassette arrays for *bla*_GES-5_ integrons, all of which were class 1 integrons^14^. In contrast, we identified 8 *bla*_GES-5_ contigs containing class 3 integron integrases, consistent with *bla*_GES-5_ transfer events between integron classes.

We then applied Flanker to the dataset of *bla*_GES-5_ contigs, revealing shared genetic structures both upstream and downstream of the gene in an exploratory analysis. Next, we constructed a pangenome graph for the dataset with PanGraph and used the shared ‘blocks’ of sequences to cluster the contigs. This approach seemed robust to both variable gene cassette synteny, and the inclusion of short or incomplete contigs. Our findings suggested different lifestyles for integrons found on the chromosome versus plasmids, where the former is well-conserved with structural variants evolving from interactions from parasitic transposable elements, and the latter is more structurally diverse. Lastly, we examined the potential of SNV profiling for integron epidemiology, first proposed in Partridge et al. 2009^3^. Overall, the utility of SNV profiling for *intI1* integrases and *bla*_GES-5_ gene cassettes remains unclear on a global scale, and perhaps better suited to localised geographies whereby invading integrons can be better distinguished from native integrons.

Our analysis has limitations. *bla*_GES-5_ gene cassettes were usually found in the first position, immediately downstream of the integrase. Yet, relying solely on sequence data meant it was impossible to determine whether this meant higher expression of the gene. Moreover, identifying a gene cassette in the first position might also indicate that it was the most recently acquired^74^, but this could not be explored here since the gene cassettes might have shuffled since a capture event. Also, by only selecting contigs with *bla*_GES-5_ combined with the fragmentary nature of integron sequencing, it is possible that our methodology favoured selecting integrons with *bla*_GES-5_ in the first position. Overall, this made it challenging to draw meaningful inferences about the evolutionary history of *bla*_GES-5_. Additionally, since we constructed our dataset from NCBI, it was susceptible to its clinical and taxonomic biases. Therefore, it might disproportionately represent studies which selectively sampled for carbapenemase-producing isolates and/or isolates from patients on cephalosporins/carbapenems, which favours the *bla*_GES-5_ phenotype and hence close proximity to the promoter. In a non-clinical sample, *bla*_GES-5_ might be more varied in position.

There were also challenges when determining the putative activity of the integrons. Our approach relied on annotations for the integrase promoters P_c_ and P_int_, and therefore hinged on a reference database. Though not feasible here, integrase functionality should be experimentally validated. This might explain that whilst both promoters were annotated within *P. aeruginosa* ST235 integrons, the order of the gene cassettes remained mostly static. Conversely, the conservation of these integrons might have been influenced by the IS*Pa21* insertions, which might disrupt *attC* sites and recombination activity. Currently, limited experimental evidence exists on the impact of IS*1111*-group insertion sequences on integron function and gene cassette expression. Furthermore, when evaluating integron function, we also did not investigate the distribution of Shine-Dalgarno sequences in the *attI* sites, which might contribute to gene cassette expression^75^.

It was also challenging to classify the contigs as plasmid or chromosomal. Many contigs were extremely short and could not be typed because the discriminating information was not present: chromosomal housekeeping genes or plasmid replicon genes. Ultimately, if the information is not present in a contig, it cannot be classified with full confidence. Also, due to chromosomal genomic islands, it is possible that some plasmid classifications were false positives. For example, in *P. aeruginosa* ST235, often contains a genomic island known as GI2, which is known to house *bla*_GES-5_^72^. Compared to acquiring a plasmid-borne ‘mobile integron’, this might be an important pathway for long-term integron acquisition within a strain.

Additionally, PanGraph was developed for complete genomes, so it is calibrated for long homologous blocks. Passing in fragmented short contigs is not its intended use, so the higher proportion of short homologous blocks could produce unexpected results. This was not investigated here but should be explored more systematically in further studies.

Though not performed here, the simultaneous analysis of the flanking sequences of other GES variants might be appropriate, considering how closely related many are (Figure S1a). Within *bla*_GES-5_ alone, we identified 12 synonymous SNV sites, which might alter phenotype, or facilitate the evolution between GES variants as ‘bridging genes’. Hence, a genomic or genetic context for one GES variant might indeed be found for another. However, incorporating this extra axis was out of the scope of this analysis.

In summary, we have presented a global analysis of *bla*_GES-5_ genetic contexts, including an approach to NCBI dataset curation, followed by exploratory and in-depth analysis. Whilst many contigs were too short to be classified as chromosomal or plasmid, a pangenome graph approach was able to give insights into the integrons involved in globally disseminating *bla*_GES-5_. Chromosomal *P. aeruginosa* ST235-associated integrons were highly conserved, whilst plasmid IncP6 integrons were diverse. This suggests that reference-based approaches are more suitable for the former, whilst flexible graph-based approaches better suit the latter. Future studies should seek to better characterise the mechanisms and dynamics driving these distinct lifestyles of *bla*_GES-5_-associated integrons.

## Acknowledgements

The authors would like to thank A. Sarah Walker, Célia Souque, and Sally R. Partridge for the insightful conversations.

---

**Figure S1.**
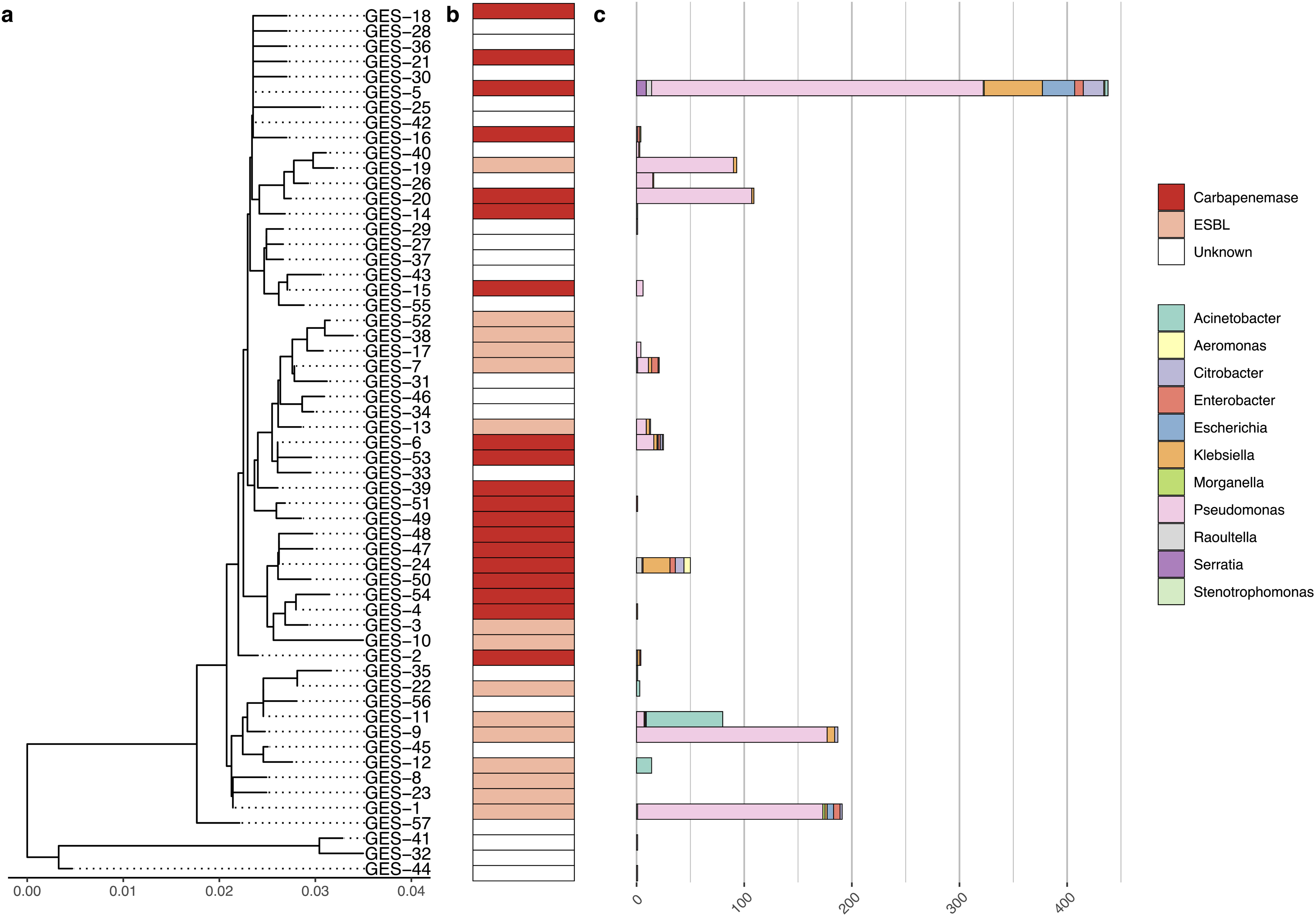
Distribution of NCBI GES variants. **(a)** Midpoint rooted phylogeny of *n*=57 GES variants, the *x*-axis represents the number of nucleotide substitutions per site . **(b)** Experimentally determined hydrolytic profile^36^. ‘Unknown’ represents missing data. **(c)** Genera distribution of NCBI contig annotations.

**Figure S2.**
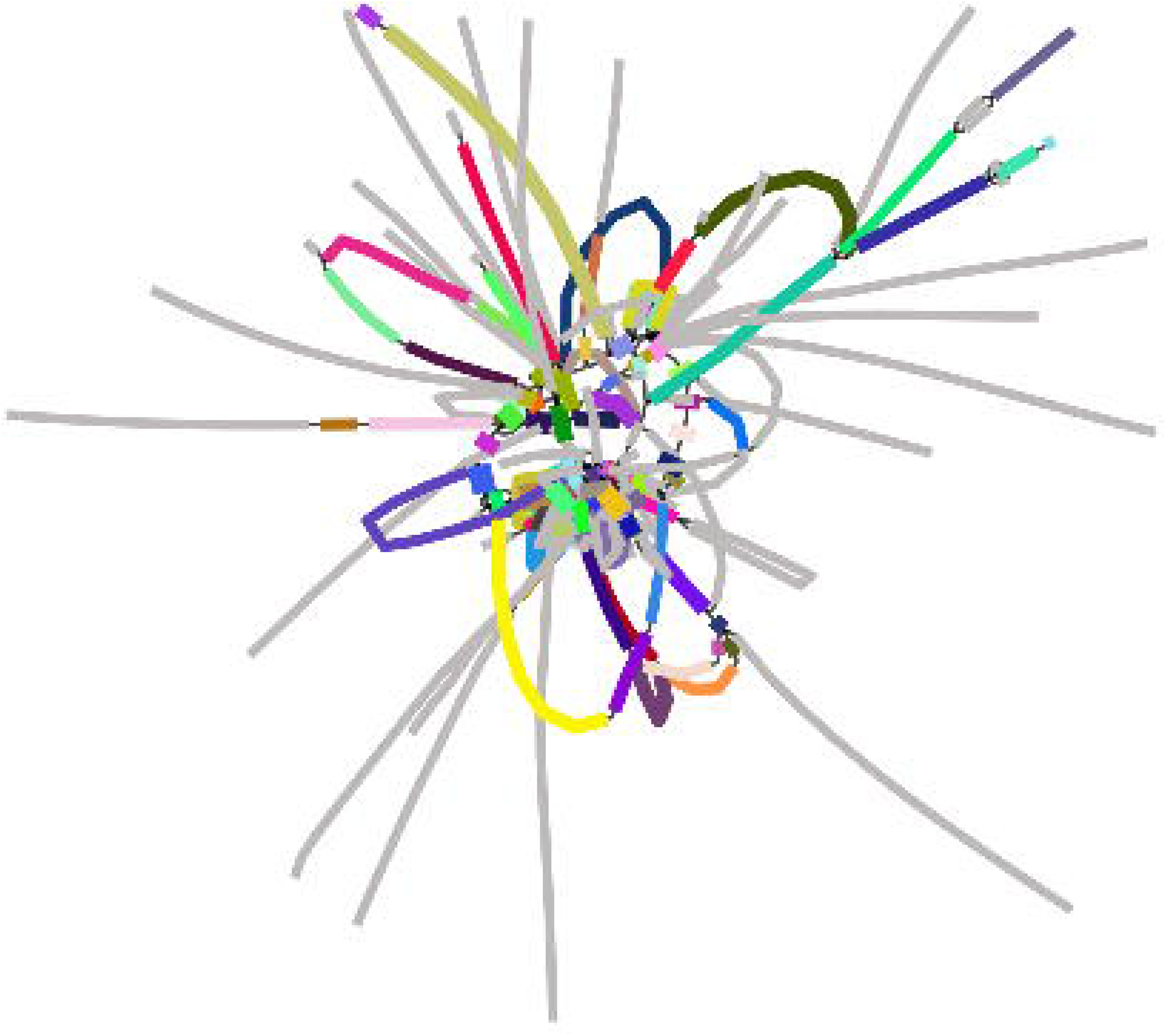
*bla*_GES-5_ 10,000bp flanking sequence PanGraph visualised in Bandage.

**Figure S3.**
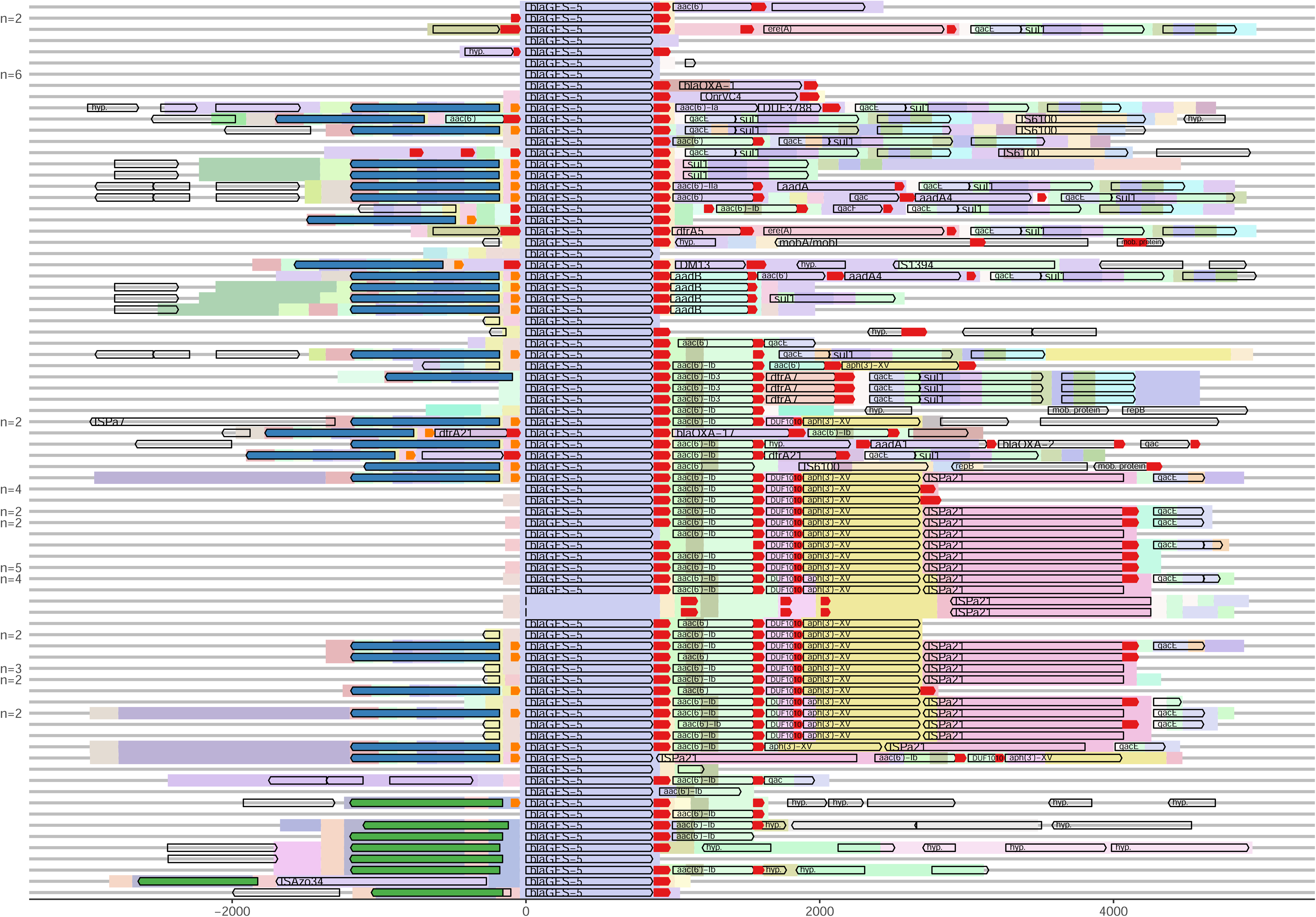
*Gene cassette annotations for* bla*_GES-5_-associated integrons.* Flanking sequences are ordered as in Figure 4, and overlayed with CDS annotations (black outlined arrows) and integron-specific annotations: *intI1* in blue, *intI3* in green, *attI* in orange, and *attC* in red. Numbers to the left indicate duplicates. Gene names are provided for any annotation found between any two *attI*/*attC* sites, as well as for 3’-conserved genes *qacE* and *sul1*.

**Figure S4.**
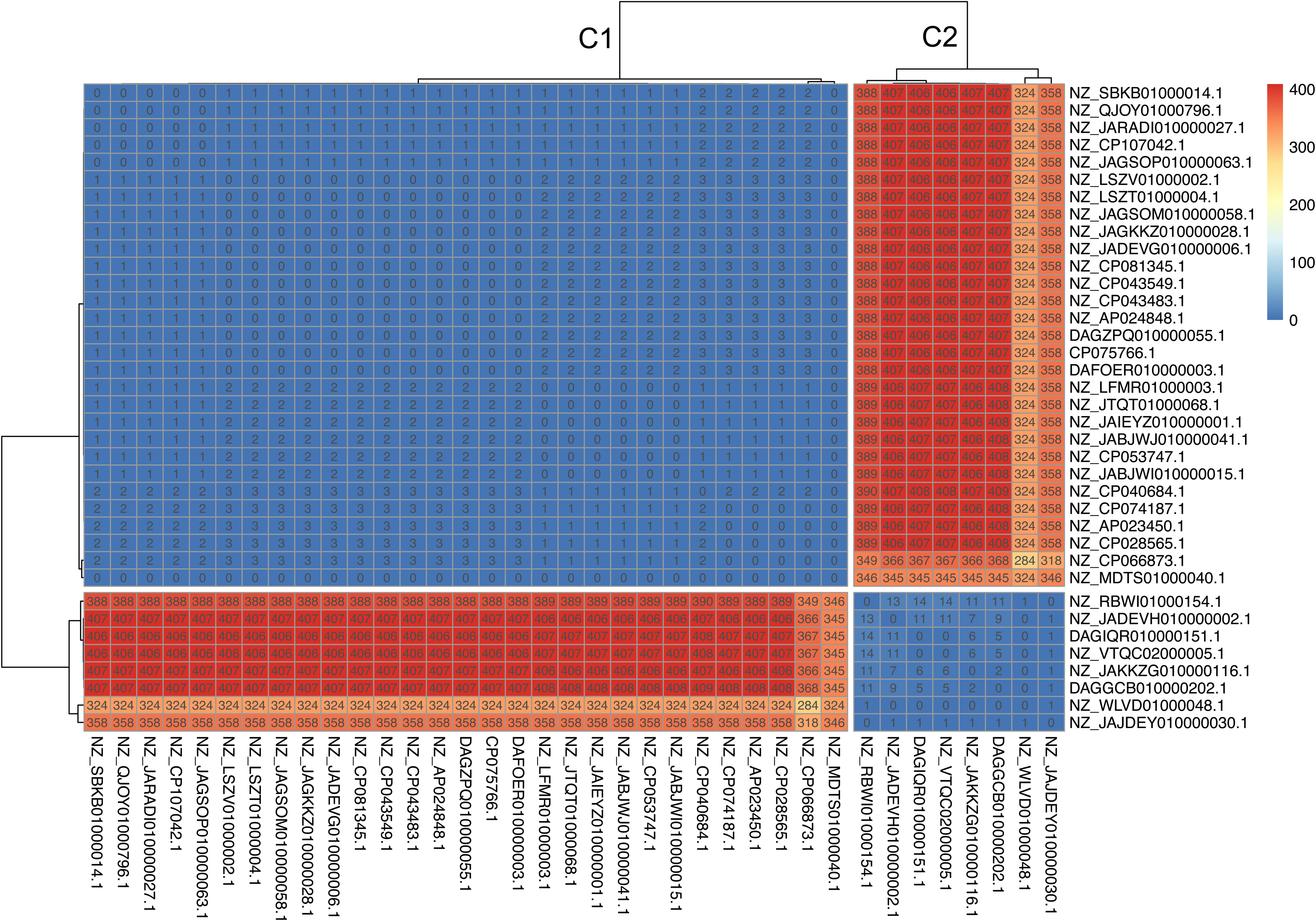
*Heatmap of integrase SNPs shows clusters for* intI1 *(C1) and* intI3 *(C2).* Numbers in cells indicate the number of SNPs between integrase pairs.

**Figure S5.**
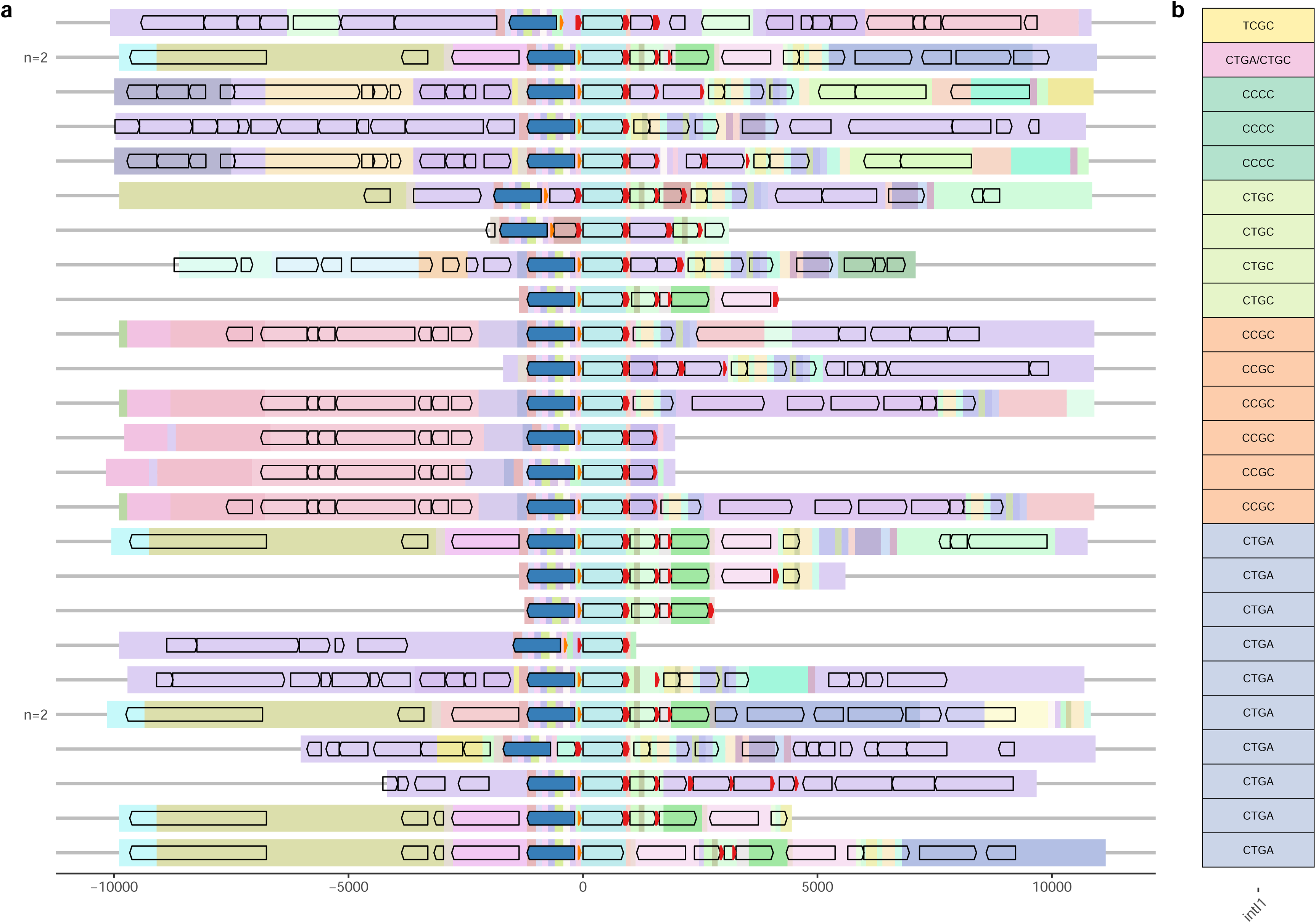
Full-length flanking sequences. **(a)** 10,000bp *bla*_GES-5_ flanking sequences containing *intI1* (blue arrow, flanking sequences subset from Figure 4) **(b)** grouped by *intI1* SNV profile.

